# Complex regulatory role of DNA methylation in caste- and age-specific expression of a termite

**DOI:** 10.1101/2022.03.08.483442

**Authors:** Mark C. Harrison, Elias Dohmen, Simon George, David Sillam-Dussès, Sarah Séité, Mireille Vasseur-Cognet

## Abstract

The reproductive castes of eusocial insects are often characterised by extreme lifespans and reproductive output, indicating an absence of the fecundity/longevity trade-off. The role of DNA methylation in the regulation of caste- and age-specific gene expression in eusocial insects is controversial. While some studies find a clear link to caste formation in honeybees and ants, others find no correlation when replication is increased across independent colonies. Although recent studies have identified transcription patterns involved in the maintenance of high reproduction throughout the long lives of queens, the role of DNA methylation in the regulation of these genes is unknown. We carried out a comparative analysis of DNA methylation in the regulation of caste-specific transcription and its importance for the regulation of fertility and longevity in queens of the higher termite, *Macrotermes natalensis*. We found evidence for significant, well-regulated changes in DNA methylation in mature compared to young queens, especially in several genes related to ageing and fecundity in mature queens. We also found a strong link between methylation and caste-specific alternative splicing. This study reveals a complex regulatory role of fat body DNA methylation both in the division of labour in termites, and during the reproductive maturation of queens.

## Introduction

DNA methylation, the epigenetic modification of DNA, is widespread among eukaryotes and is known to be important for transcriptional regulation of genes and repression of transposable elements (Zemach et al., 2010). Age-related changes in DNA methylation levels and an increased variability known as epigenetic drift have been recognised as an important hallmark of ageing in mammals (Issa et al., 2014; López-Otín et al., 2013). DNA methylation has garnered considerable attention within social insects with an apparent role in the regulation of sterile and fertile castes in honey bees (Lyko et al., 2010) and in ants (Bonasio et al., 2012). A more recent study found a significant role of methylation in the task division of worker bees (de Souza Araujo and Arias, 2021). However, there remains considerable debate surrounding the universality of the role of DNA methylation in the transcriptional regulation of caste-specific genes in eusocial insects (Herb et al., 2012; Patalano et al., 2015; Libbrecht et al., 2016). In bumble bees, DNA methylation appears to be more important for worker reproduction (Amarasinghe et al., 2014) than for caste differentiation (Marshall et al., 2019). Two studies found no influence of DNA methylation on the formation of behavioural castes in a wasp (Patalano et al., 2015) and an ant (Patalano et al., 2015; Libbrecht et al., 2016) that live in simple societies. In fact, the authors of the latter study claimed previous evidence for the role of DNA methylation in the division of labour was weak and that further studies required more robust methodology, especially greater replication (Libbrecht et al., 2016). Most of these studies have concentrated on social Hymenoptera (ants, bees and wasps), with the exception of two studies on the role of DNA methylation in the division of labour in adult termites. The first of these studies investigated whole-body methylation patterns for the lower, drywood termite *Zootermopsis nevadensis* (Glastad et al., 2016), which forms simple colonies, in which workers retain the possibility to become fertile (Weil et al., 2007). In the second study, head methylomes of the subterranean termite, *Reticulitermes speratus*, were investigated, a species with an intermediate level of social complexity (Shigenobu et al., 2022). While the first study found large differences between castes in *Z. nevadensis* (Glastad et al., 2016), Shigenobu et al. (2022) found very strong correlations in DNA methylation patterns between castes of *R. speratus*. However, in the first study, limited replication was performed within one single colony, while in the second study non-replicated castes were sampled from different colonies, so that the effect of colony-specific variation, inherent in previous studies (Libbrecht et al., 2016), could not be excluded in either of these studies. The general role of DNA methylation in the transcriptional regulation of termite castes is therefore still unclear, especially in higher termites that form complex colonies with lifelong sterile worker castes.

Beside reproductive division of labour, the eusocial insects are also characterised by extreme longevity among fertile castes, indicating an apparent absence of the fecundity-longevity trade-off attributed to non-social insects (Korb et al., 2021). Several, recent studies have presented evidence for the transcriptional regulation of specific gene co-expression modules associated with old but highly fertile queens in ants (Harrison et al., 2021), bees (Séguret et al., 2021) and termites (Lin et al., 2021; Séité et al., 2022). However, the role of DNA methylation in this absence of the longevity-fecundity trade-off in eusocial insects is so far unknown.

In this study, we investigated caste- and age-specific DNA methylation profiles to make inferences on the regulation of genes important for the extreme longevity and high fecundity of reproductives in the higher termite, *Macrotermes natalensis*. This foraging, fungus-farming termite is characterised by large colonies and sterile workers. Kings and queens can live for over 20 years (Keller, 1998), with the highly fertile queen laying thousands of eggs per day (Kaib et al., 2001). The mature *Macrotermes* queens are characterised by a hypertrophic abdomen, as well as several further metabolic and physiological differences compared to virgin queens, such as enlarged corpora allata (Sieber and Leuthold, 1982), increased DNA content and major changes in insulin signalling and fat storage (Séité et al., 2022).

We carried out reduced representation bisulfite sequencing (RRBS) on four phenotypes (short-lived, sterile female workers, young virgin queens, 20-year-old queens, and 20-year-old kings), replicated across three independent colonies from this higher termite and related DNA methylation patterns to caste- and age-specific gene expression. This was performed on the fat body, since we recently showed the importance of this tissue for the long reproductive life of the reproductive termite castes (Séité et al., 2022).

## Results and Discussion

### RRBS is a robust method for determining genomic methylation patterns in termites

For each of the four phenotypes, female workers (FW), virgin queens (VQ), mature queens (MQ) and mature kings (MK), we aimed to produce reduced representation bisulfite sequencing (RRBS) for 3 replicates from independent colonies. An accurate estimation of methylation levels relies heavily on an efficient conversion rate of unmethylated sites with the bisulfite treatment. To measure the erroneous, non-conversion rates, each sample was supplemented with a non-methylated lambda spike-in control (see methods). We kept only those samples with a non-conversion rate lower than 2% (Table S1). We generated between 32.1M and 61.4M bisulfite treated reads per sample (Table S1). These reads were mapped to the genome (mapping rate: 67.3%-71.2%; Table S1) to quantify methylation levels, and for each sample only CpGs to which at least 5 reads mapped were included in analyses. We were able to quantify methylation levels (at least 5 reads) of 6.29 million CpG sites (19.1% of all genomic CpGs). For each phenotype most CpGs were sequenced for all 3 replicates, ranging from 2.8M to 3.6M CpGs per phenotype (Fig. S1A-D). In support for the reliability of the RRBS method, a large proportion of the CpGs (1.97M, 31.3%) were sequenced consistently within all 12 samples (4 phenotypes x 3 replicates), which was by far the largest intersection of the 12 sets of sequenced CpGs (Fig. S1E). All subsequent analyses are based on this subset of 1.97M CpGs.

### High gene body methylation

Within the subset of 1.97M CpGs that were sequenced within all 12 individuals, we found detectable methylation at 49.0% (FDR corrected binomial p-value < 0.05, based on non-conversion rate) of sites in at least one sample. For each of the 12 samples, methylation level was calculated for each sequenced CpG as the proportion of mapped reads that were putatively methylated (non-converted cytosines). To estimate overall genomic methylation levels, we calculated means across the 12 samples at each CpG. Methylation levels varied throughout the genome, with highest rates within coding regions (mean 9.74% per CpG, standard error: 0.20) and lowest rates within intergenic regions (mean: 1.70%, SE: 0.01; Fig. 1A). In repetitive regions, methylation was higher than in intergenic regions (mean: 2.22%, SE: 0.01), indicating that transposable elements (TEs) may be targeted by DNA methylation. Similar to findings for the lower termite, *Z. nevadensis* (Glastad et al., 2016), methylation was relatively high in introns (mean: 3.91%, SE: 0.05 Fig. 1). In support of findings for *Z. nevadensis* (Glastad et al., 2016) but in contrast to Hymenoptera (Bonasio et al., 2012; Patalano et al., 2015), we found that, for all samples, methylation levels increased along the gene body, with highest levels at 3’ exons (13.6-23.3% among 5th to last exons) and introns (10.4-16.8% among 4th to last introns; Fig. 1B), suggesting this gene body methylation pattern may be widespread among termites.

**Figure 1:**
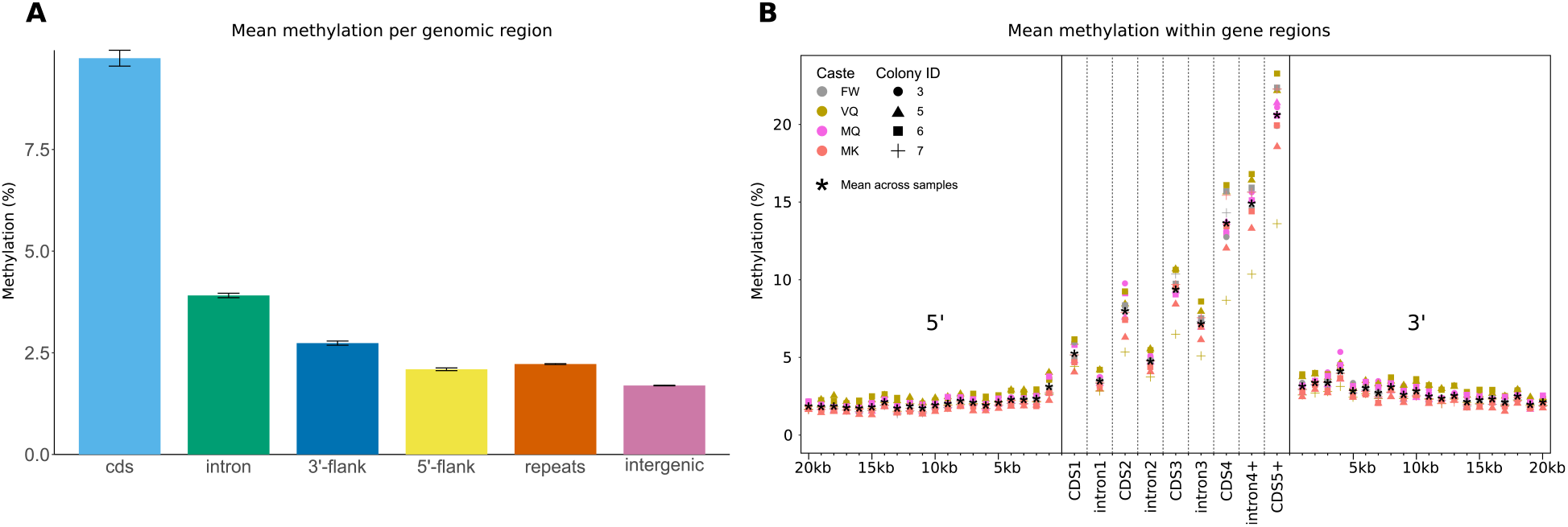
Genomic variation in methylation. In **A.** mean methylation proportions among all 12 samples are shown for 6 categories of genome regions. Error bars are standard error. Flanks are defined as 10kb up- or downstream of coding regions. **B.** Mean methylation (%) within gene bodies (exons and introns) and in twenty 1kb bins at 5’- and 3’-flanking regions of genes. Each dot represents mean methylation for one of twelve samples across all sequenced CpGs within the region of interest. The four phenotypes (FW, VQ, MQ, MK) are represented by colour; the colonies, from which replicates originated, are represented by shape. Stars show means across all 12 samples.

### Greater variation in methylation between colonies than between phenotypes

We detected high individual variation in methylation patterns, with 24.7% to 26.2% of CpGs methylated in only 1 of the 12 samples, while only 9.1% to 9.7% were methylated in 2 individuals. Interestingly, as previously found in the clonal raider ant, *Dinoponera quadriceps*(Libbrecht et al., 2016), we found a substantial number of CpGs (8.4%) within coding sequence and introns (2.7%) that were robustly methylated within all 12 samples (Fig. 2A). These robustly methylated CpGs were situated in genes enriched for GO-terms related to cell differentiation, cell adhesion and regulation of cellular processes (Table S2). Interestingly, robustly methylated genes (containing at least one CpG methylated in all 12 samples) were more frequently differentially expressed between phenotypes (89.6%), compared to other genes (64.9%), suggesting an important role of DNA methylation in the regulation of gene transcription. Furthermore, methylation patterns (proportion of methylated reads per CpG), correlated strongly and uniformly between all samples (Pearson’s r: 0.600-0.781; p-value = 0), especially within coding sequence (0.889-0.960), indicating little differentiation between castes, similar to findings for the subterranean termite, *R. speratus* (Shigenobu et al., 2022). The slightly lower correlations we report here compared to those found for *R. speratus* may be linked to a number of differences in this current study, such as colony replication, RRBS rather than whole genome BS-sequencing, or may be related to species-specific patterns. Furthermore, the high correlations we found between VQ and MQ (0.663-0.778) suggest DNA methylation patterns are well maintained with age in termite queens. This apparent lack of epigenetic drift, at least for DNA methylation, may help to explain the recently documented, well-regulated transcription of anti-ageing genes in *M. natalensis* queens (Séité et al., 2022).

**Figure 2:**
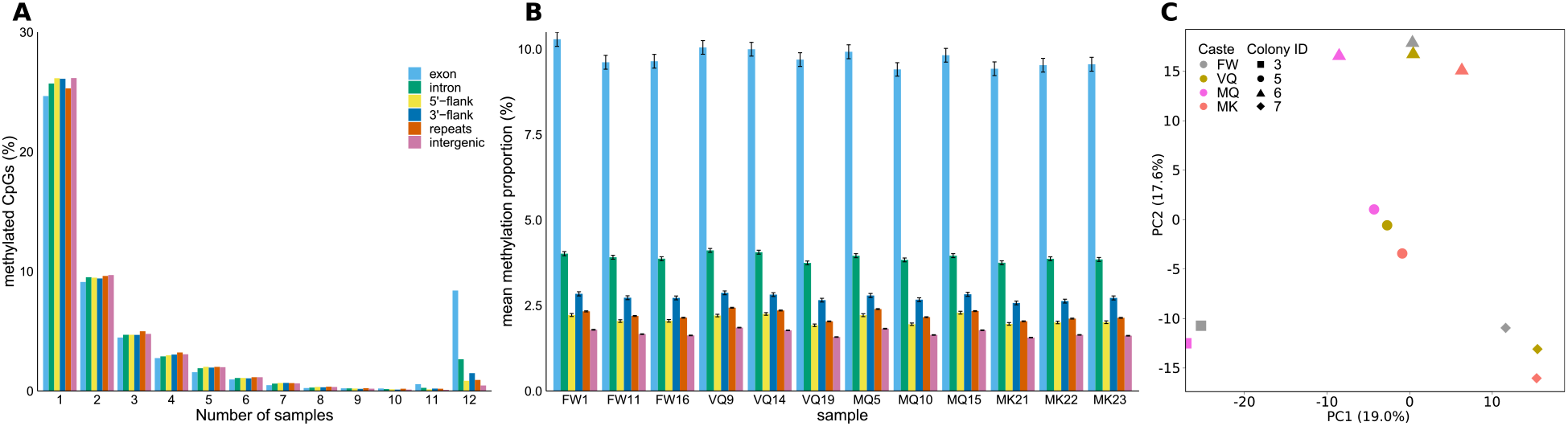
Individual variation in methylation. **A.** Proportions of CpGs that are methylated (FDR < 0.05) in varying numbers of 12 samples within 6 genomic regions. **B.** Mean proportions of methylated reads across all CpGs for each of the 12 samples (4 phenotypes x 3 replicates). **C.** Principal component analysis of methylation at 1000 most variable CpGs in 12 samples, spanning four phenotypes, represented by colour (FW, VQ, MQ and MK), from four colonies, represented by shape. The first two principal components are displayed on the x- and y-axes with variance explained in brackets.

Methylation levels (proportion of methylated reads) also varied among individuals, with coding methylation ranging from mean 9.41% (± 0.20 standard error) in the mature queen from colony 5 (sample ID: MQ10) to 10.29% (± 0.21 SE) in the female worker sample from colony 3 (FW1; Fig. 2B). Intergenic CpGs, on the other hand, were most highly methylated in the VQ sample from colony 5 (VQ9; mean: 1.85% ±0.01 SE) and lowest in the MK sample from colony 5 (MK21; mean: 1.56% ±0.01 SE). A principal component analysis revealed that methylation patterns vary more between colonies than between castes (Fig. 2C) as previously found for the ants, *Cerapachys biroi* (Libbrecht et al., 2016) and *Dinoponera quadriceps*, and the paper wasp, *Polistes canadensis* (Patalano et al., 2015). This highlights the importance of replication across independent colonies in methylation studies as previously reported (Libbrecht et al., 2016), thus raising the question of whether caste-specific methylation patterns detected within a single colony for the lower termite *Z. nevadensis* were species- or colony-specific (Glastad et al., 2016). High colony variation is confirmed by a 3-way ANOVA among the 10 000 most variable sites, in which colony (F(3,1.20×10^5^)=843.2, p = 0.0) has an effect size (generalised eta squared[ges]=0.021) larger than that of phenotype (F(3,1.20×10^5^)=492.1, p = 7.70×10^−318^, ges=0.012), while genomic region (exon, intron, 5’-flank, 3’-flank, repeats, intergenic) was an even stronger predictor of methylation level (F(5,1.20×10^5^)=720.0, p = 0, ges = 0.029). However, significant interactions existed between all three factors, indicating differing effects of each combination of phenotype, colony membership and genomic region on methylation level.

### Conserved, single-copy genes are more highly methylated

We performed two analyses which confirmed higher methylation levels for conserved genes. We first analysed gene age by determining the broadest phylogenetic taxon for which a gene ortholog could be found, ranging from species-specific to Mandibulata. The proportion of highly conserved genes, found in the oldest category, Mandibulata, was highest among genes with methylation levels greater than 80%, while species-specific genes were proportionally most abundant among lowly methylated genes (Fig. 3A). In further support for greater methylation of conserved genes, we found significantly higher methylation levels among single-copy ortholog genes (single copy in *M. natalensis* with orthology in other insects) than in multi-copy genes (> 2 paralogs). Similarly, for singletons (single-copy, species-specific genes), which are likely evolutionarily novel compared to orthologs, methylation levels were lower than in single-copy orthologs and did not differ from multi-copy genes. The methylation of 2-copy genes were intermediate between single-copy and multi-copy genes (Fig. 3B).

**Figure 3:**
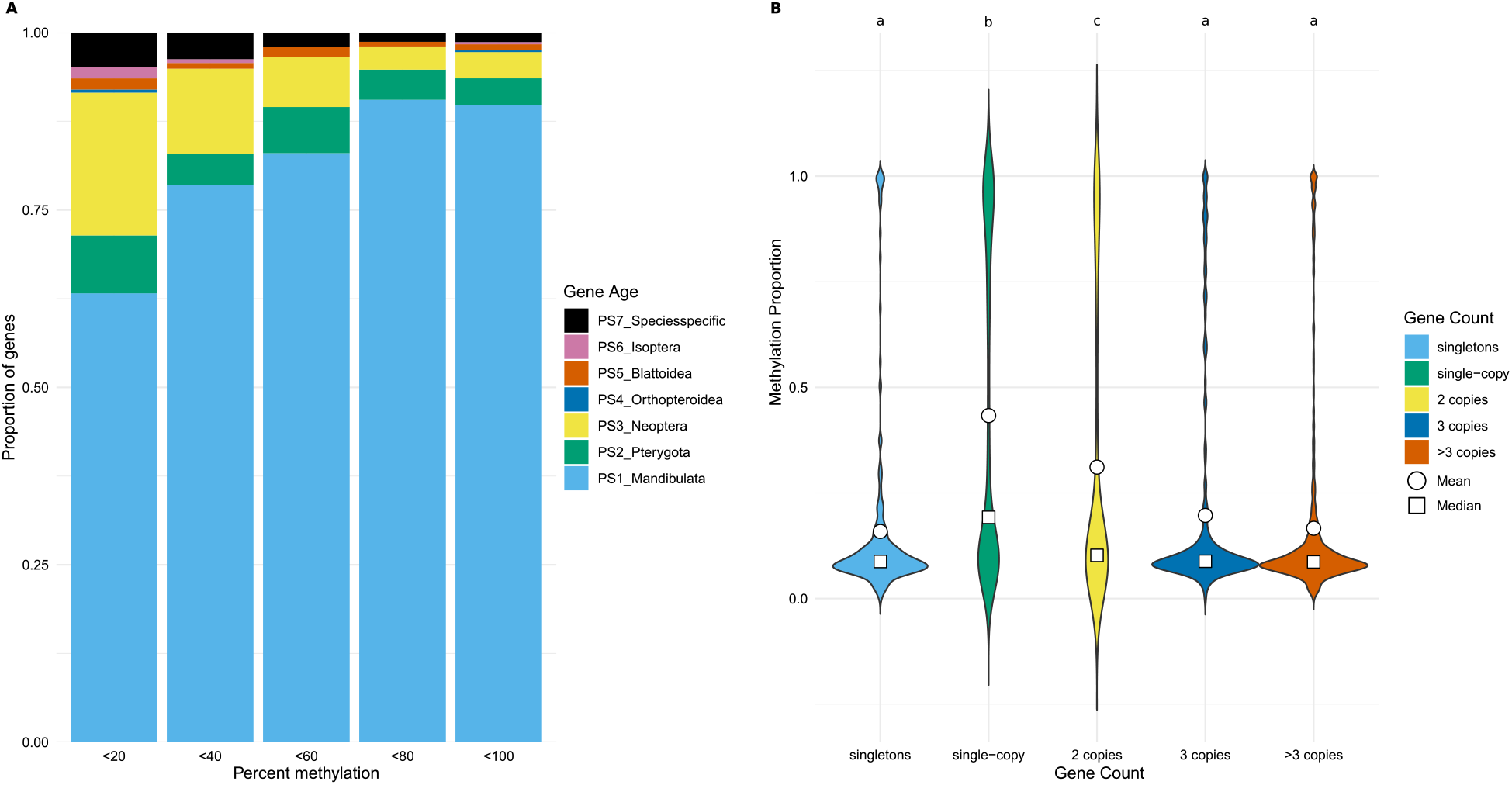
Methylation and gene conservation. A. Proportions of gene age categories within 5 categories of methylation level. B. Methylation level within genes with varying numbers of copies. Singleton = no paralogs or orthologs; single-copy = no paralogs but with orthologs in other species; other gene groups have varying numbers of paralogs.

### Ageing and fertility genes hypomethylated in mature queens

Despite the larger variation between colonies, we found 1344 CpG sites to be significantly differentially methylated (DMS) between phenotypes. We tested whether these numbers of DMS are greater or smaller than can be expected between two groups of three randomly assigned samples (1000 bootstraps; 95% confidence interval: [45-102]; 99% confidence interval: [40-114]). In this manner, we found a significant number of DMS that were hypermethylated in VQ compared to each of the other castes (> 95%). In MQ, on the other hand, there were significant numbers of DMS that were hypomethylated compared to other castes (> 95%; Fig. 4A). The numbers of unique DMS varied among phenotypes and genomic regions, and were enriched within coding regions (2.7-5.0%) compared to the proportion of total sequenced CpGs within coding regions (1.2%). Interestingly, the largest category of DMS were those hypomethylated in MQ (365 unique sites) while the smallest category contained sites hypomethylated in VQ (163) (Fig. 4B). Of the 1291 DMS, 386 lay within 261 genes (DMGs), of which 111 genes contained sites hypomethylated in MQ, while 87 genes contained sites hypermethylated in VQ. These striking results indicate a major shift in methylation patterns occurs during queen maturation for a subset of genes.

**Figure 4:**
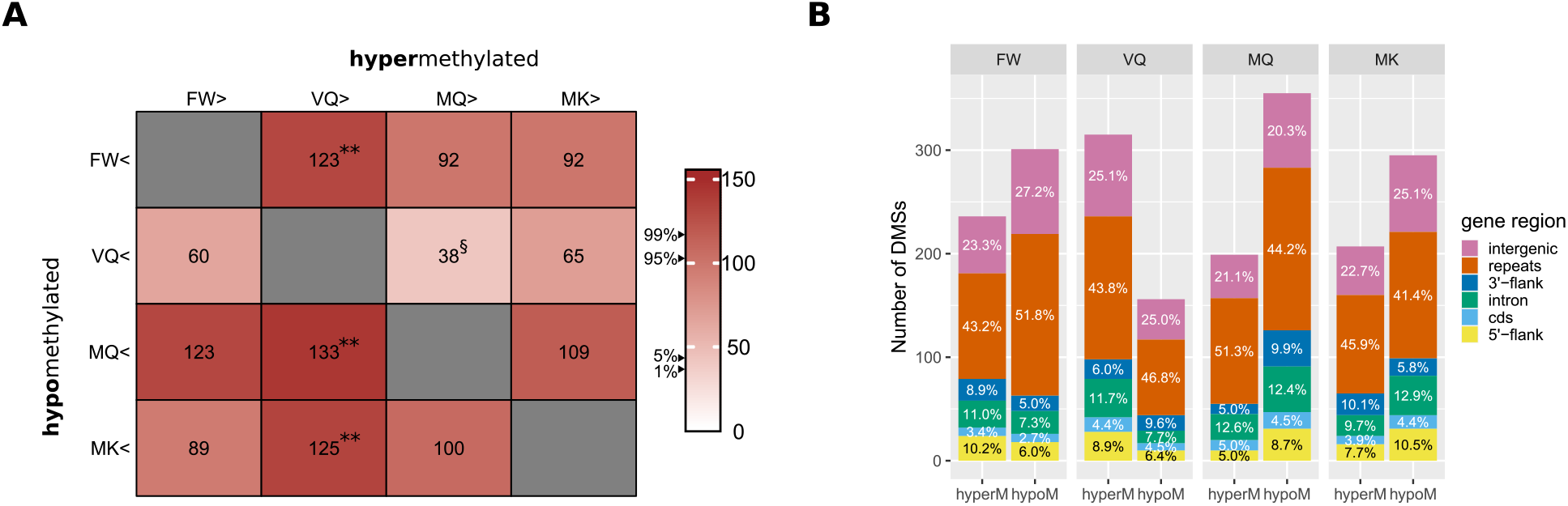
Differentially methylated sites. **A.** Numbers of CpG sites hyper- (columns) and hypomethylated (rows) between pairs of phenotypes. Bootstrapping was carried out based on numbers of significant sites in 1000 comparisons between randomised 3×3 samples; 95% confidence interval: [45-102]; 99% confidence interval: [40-114]. ** > 0.99; * > 0.95; § < 0.05. **B.** Proportions of DMSs per genomic region for each phenotype. Unique DMSs were counted from all pairwise comparisons between the four phenotypes.

Several of the genes with significantly decreased methylation in MQ compared to VQ have important roles in ageing, including 2 regulators of Notch signalling, 2 genes involved in Wnt signalling, a Sirtuin, a sphingomyelinase, important for cellular stress, and a gene responsible for the regulation of misfolded proteins (Table S3). Further genes are related to fertility such as Vitellogenin and an ecdysone receptor (Table S3). In a previous study on this species, the major importance of insulin signalling in the fat body during the maturation process of queens was highlighted (Séité et al., 2022). It is therefore striking that *chico*, the substrate of insulin receptors in the insulin signalling pathway, and *daw*, with known functions in insulin regulation, are hypomethylated and differentially expressed in MQ compared to VQ (Table S3). A large proportion of the 44 genes containing sites hypomethylated in MQ compared to VQ, were also differentially expressed: 6 were over-expressed in MQ (13.6%), 14 genes were lower expressed in MQ (31.8%) compared to VQ, while 24 (54.5%) did not differ in expression. These proportions of differentially expressed genes are significantly higher than those found in all genes (10.6% and 15.8%, respectively; χ^2^: 9.64, df = 2, p-value = 0.008), indicating an important role of DNA methylation in the regulation of age-specific expression.

Furthermore, we found that differentially expressed genes (DEGs: significantly up- or down-regulated between pairs of phenotypes) had unique, phenotype-independent methylation signatures (Fig. 5). For instance, while the full set of DEGs have a mean methylation level of 7.4% in coding regions, genes with over-expression in MQ or MK compared to VQ, or in MQ versus FW, have very low coding region methylation (2.3, 3.2% and 4.0%, respectively). Genes overexpressed in MQ and MK compared to VQ also have high methylation in 3’-flanks (3.5 and 3.1%, respectively), compared to all DEGs (2.2%) (Fig. 5). Surprisingly, within each of these DEG groups, variation among phenotypes was low, with standard deviation among samples ranging from 0.10 to 0.53. These patterns point towards a complex relationship between DNA methylation and caste- or age-specific gene expression in *M. natalensis*.

**Figure 5:**
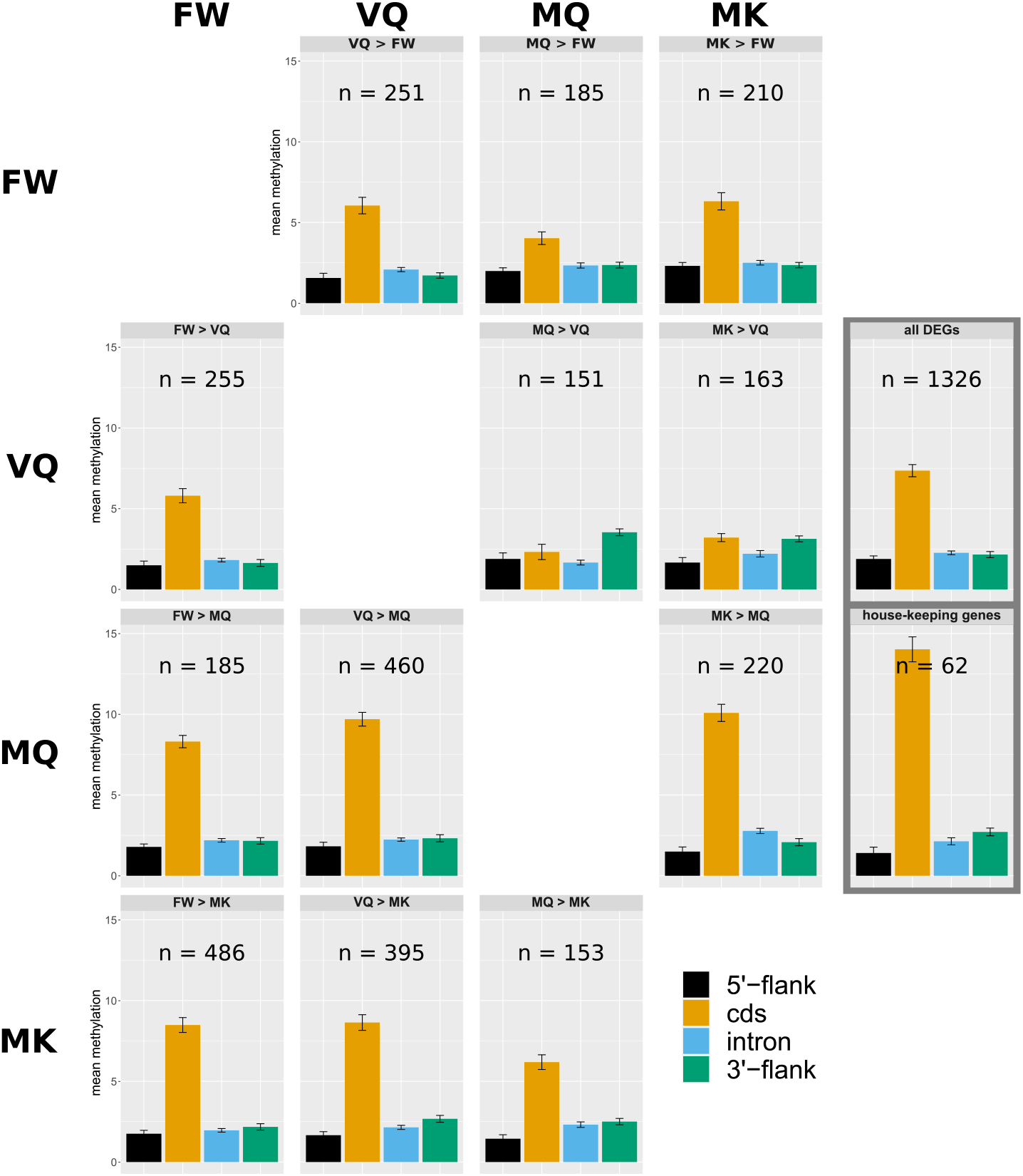
Mean methylation level per gene region for groups of differentially expressed genes. HKG = house-keeping genes, defined as non-differentially expressed genes, with expression counts greater than median expression.

### Variation in gene body methylation influenced by expression level, caste-specific expression and alternative splicing

To better understand the variation in methylation levels among genes, we first investigated the influence of expression level. We found a significant positive correlation between methylation level of coding sites and expression level, which ranged from 0.208 (p-value = 2.0×10^−176^) to 0.254 (p-value = 6.9×10^−265^; spearman’s rank correlation) per sample. This confirms previous findings for Hymenoptera (Bonasio et al., 2012; Patalano et al., 2015; Libbrecht et al., 2016) and a termite (Glastad et al., 2016). Among genes whose expression differed significantly among phenotypes (DEGs), we found a significant positive interaction with expression, with a linear regression predicting higher methylation for DEGs compared to nonDEGs for expression levels greater than the 4th percentile (Fig. 6A). We also found that methylation level increases with the number of isoforms per gene, when controlling for expression level, with methylation level predicted to be higher for multiple isoform genes at expression levels greater than the 17th percentile (Fig. 6B). For genes which are putatively differentially spliced among phenotypes (significant differential exon expression), a linear regression predicts significantly higher methylation regardless of expression level (Fig. 6C). These results suggest an important role of DNA methylation in the regulation of gene expression level, especially when regulating caste- and age-specific transcription and splicing. The regulation of caste-specific splicing via DNA methylation may be universal in eusocial insects since similar evidence has been found in honeybees (Lyko et al., 2010), ants (Bonasio et al., 2012; Libbrecht et al., 2016), and the lower termite, *Z. nevadensis* (Glastad et al., 2016).

**Figure 6:**
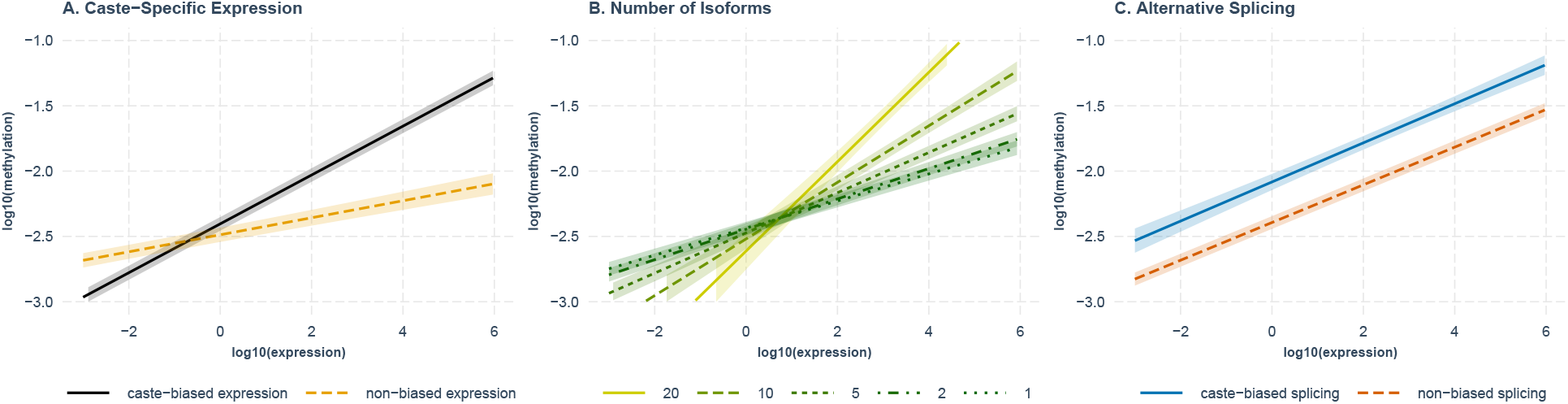
Linear models, relating differential gene and isoform expression to methylation level. **A.** Differentially expressed genes are more highly methylated when accounting for expression level. **B.** Methylation increases with increasing number of isoforms, relative to expression level. **C.** Genes that show age- and caste-specific alternative splicing are more highly methylated, regardless of expression. Models have the form: log_10_(methylation level)~ *log*_10_(expression level) * variable + (1|sample).

## Conclusions

We report a strong correlation of DNA methylation patterns with caste- and age-specific gene expression and alternative splicing in the fat body of the higher termite, *M. natalensis*. These results offer further support for the importance of fat body transcription (Séité et al., 2022) and its regulation for the extreme longevity and fecundity of termite queens. We also confirm the importance of replication in methylation analyses due to higher variation in methylation between colonies than between castes, a point of contention among previous studies in Hymenoptera (Libbrecht et al., 2016). Furthermore, and importantly, we present evidence for unique methylation signatures which are stable between phenotypes but differ especially between groups of genes with age-biased expression. For example, genes with higher expression in mature reproductives (MQ and MK) than in young reproductives (VQ) have relatively low coding region methylation but high methylation in 3’-flanks among all phenotypes compared to other DEGs. We believe this is the first time such a methylation pattern has been presented for social insects and suggests its generality should be tested on further species. We show for the first time, how DNA methylation may be responsible for regulating genes which are central to termite queens maintaining high fertility at extreme ages. For the 20-year old, highly fertile queens, we present evidence for well-maintained DNA methylation, in support of an apparent lack of epigenetic drift, a well established hallmark of ageing (López-Otín et al., 2013). Several genes with important roles in ageing and fertility, on the other hand, contain sites with significantly reduced methylation levels in mature queens compared to young, virgin queens, many of which have significantly different expression levels in old compared to young queens.

## Methods

### DNA extractions and sequencing

Total genomic DNA from the 12 termite samples (female workers, young virgin queens, mature queens and kings; see Table S1 and Séité et al. 2022 for sampling) was extracted from fat body using DNeasy Blood and Tissue kit (Qiagen), including RNase A treatment (Qiagen), according to the manufacturer’s instructions. Library construction was performed using the Premium Reduced Representation Bisulfite Sequencing kit (Diagenode). Briefly, for each sample, 100 ng of genomic DNA were digested using MspI for 12 hours at 37°C. DNA ends were repaired and Diagenode indexed adaptors were ligated to each end of the repaired DNA. Each ligated DNA was quantified by qPCR using the Kapa Library quantification kit (Kapabiosystems) on a LightCycler 480 (Roche Life Science) prior to pooling (4, 5 or 6 samples per pool). Each pool was subjected to bisulfite conversion and desalted. Optimal PCR cycle number was determined by qPCR (Kapa Library quantification kit, Kapabiosystems) before the final enrichment PCR. Once purified using magnetic beads (AMPure XP, Beckman Coulter), library pools were verified on Fragment Analyzer (AATI) and precisely quantified by qPCR using the Kapa Library quantification kit (Kapabiosystems). Each pool was denatured, diluted and spiked with a 10% phiX Illumina library before clustering. Clustering and sequencing were performed in single read 100 nt, 1 lane per pool, according to the manufacturer’s instructions on a Hiseq2500 using Rapid V2 clustering and SBS reagents. Base calling was performed using the Real-Time Analysis Software and demultiplexing was performed using the bcl2fastq software, both from Illumina. Non-conversion rate of bisulfite treatment was estimated with a spike-in control, and only samples with a non-conversion rate lower than 5% were kept for further analysis.

### Preparation of RRBS data

The RRBS reads were prepared by following the Bismark protocol (Krueger and Andrews, 2011). This included adapter trimming with Trim Galore, v.0.4.4_dev (https://github.com/FelixKrueger/TrimGalore) at default settings with the additional –rrbs argument. Subsequently, Bismark was used to analyse methylation states. The *M. natalensis* genome (Poulsen et al., 2014) was indexed using the bismark_genome_preparation command, then sequenced reads were mapped to the genome using bowtie2, version 2.3.4.3 (Langmead and Salzberg, 2012). Otherwise, standard parameters were implemented for the Bismark pipeline.

### Methylation analyses

We extracted methylation and read coverage information from the thus produced bam files with the bismark_methylation_extraotor command, with the arguments –scaffolds and –bedGraph. We only considered sites to which at least 5 reads mapped. Based on the non-conversion rate of a spike-in control, a binomial test was carried out to confirm the significance of a measured proportion of non-converted, and therefore putatively methylated, reads, as previously performed by Glastad *et al*. (Glastad et al., 2016). P-values were FDR corrected, and only corrected p-values < 0.05 were deemed methylated, and were otherwise counted as non-methylated. Sequenced cytosines (≥ 5 reads) were annotated with gene features - exons, introns, 10kb flanking regions, repetitive regions - based on information stored in two GFF files, containing protein coding and repeat element annotations (Harrison et al., 2018; Poulsen et al., 2014).

### Principal component analysis (PCA)

The PCA analysis was performed in R, version 4.0.2 (R Core Team, 2016). For each CpG site that was covered by at least 5 reads in all 12 samples, we measured variance in methylation among samples and selected the 1000 most variable sites. The PCA was computed on these top variable sites with the prcomp function and the first two PCs were plotted with ggplot2 (Wickham et al., 2016).

### Regression models

For each gene, average methylation level was calculated per feature type (exons, introns, 5’-flank and 3’-flank) and per sample. All regression analyses were performed on this data set. The following variables were considered:

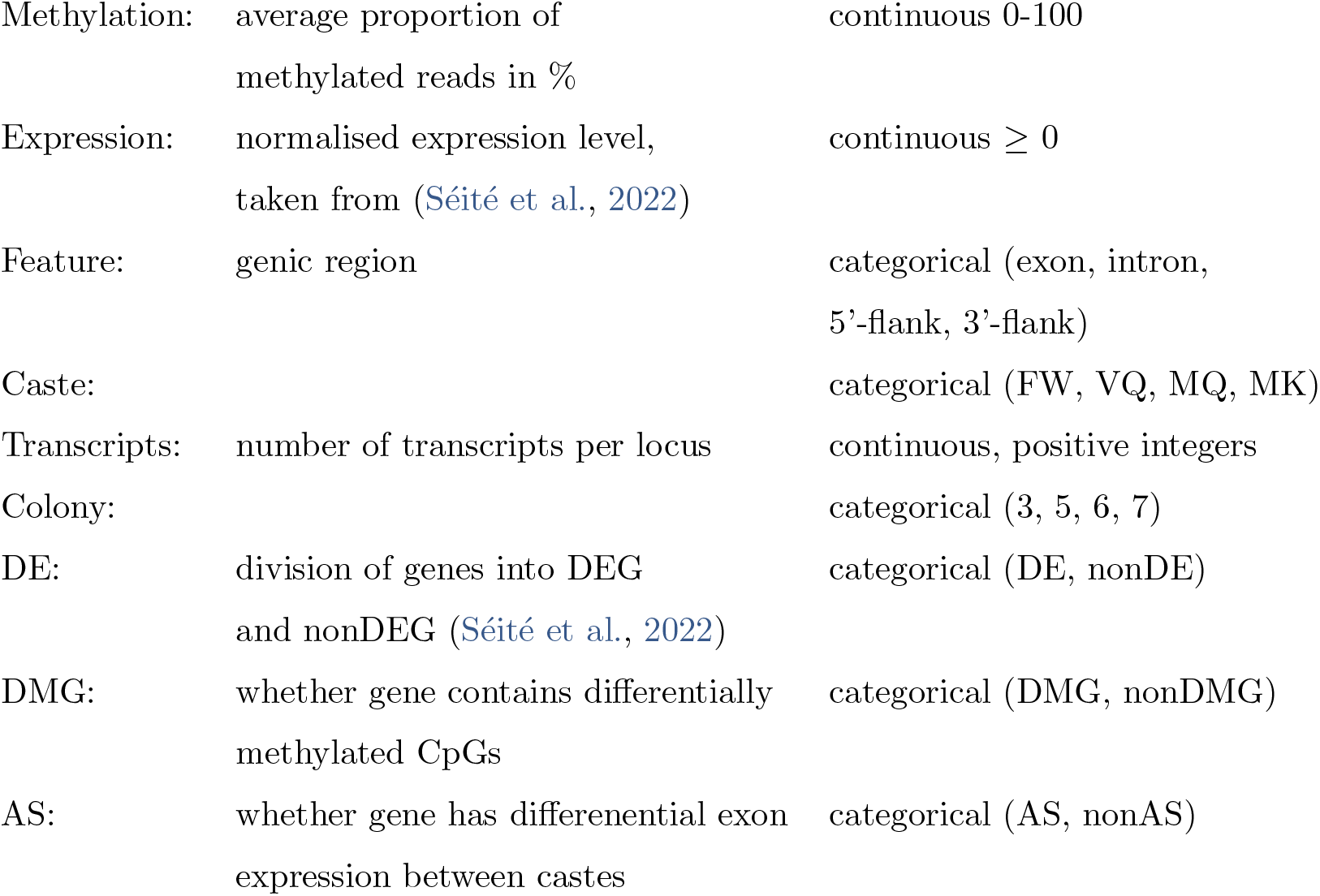

ANOVAs and ANCOVAs were also performed in R with the anova_test function from the rstatix library (Kassambara, 2021). For graphical representations, we used the lmer function from the lme4 package (Bates et al., 2015) to create the model and interact_plot from the interactions package (Long, 2019) for plotting. In each case, we log-transformed expression and modelled non-liner regression of methylation with the poly function, using as many polynomials as were significant. The following variables were included as co-factors: number of transcripts per gene, caste membership, genomic feature (exon, intron, 5’-flank, 3’-flank), differential expression; with colony membership as the random effects term. For example, to relate methylation to expression by caste and differential expression, while controlling for colony membership:

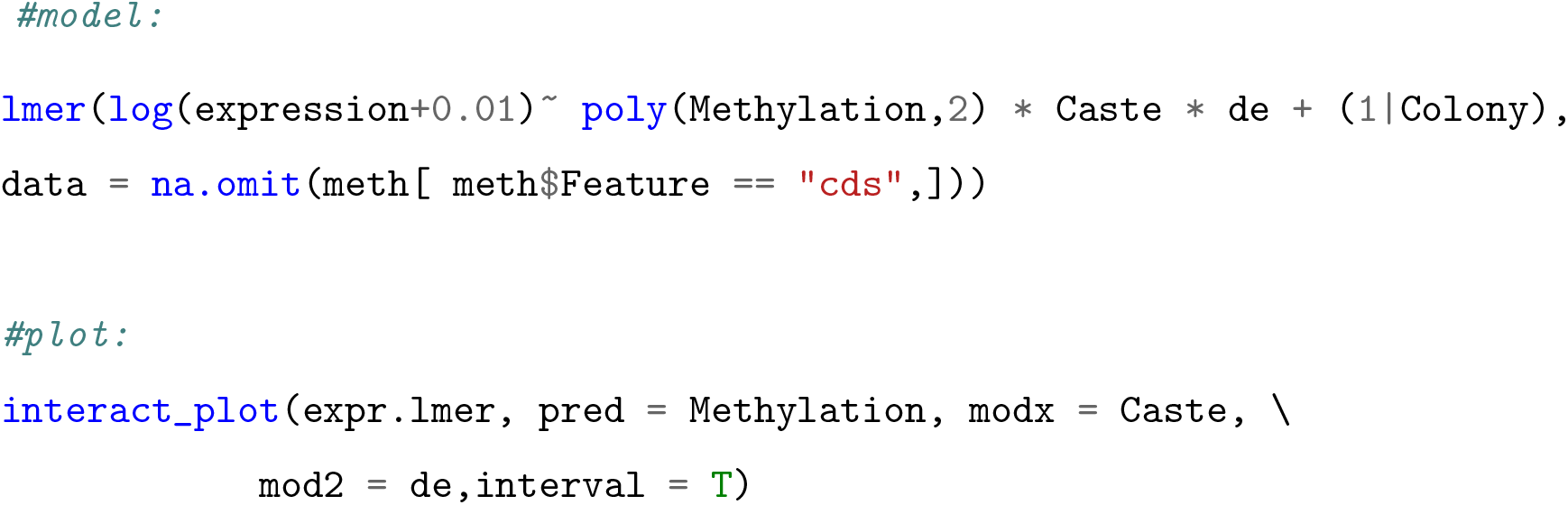

### Detecting differential methylation

To detect significant differences in methylation between phenotypes, we used the R package methylKit, version 1.11.1 (Akalin et al., 2012). We analysed differential methylation between all pairs of the four phenotypes (FW, VQ, MQ, MK) and for each of these comparisons only included CpGs, for which at least 10 reads existed for all 6 samples (3 replicates x 2 phenotypes). A difference in methylation was only considered significant if it were at least 25 percentage points and with an adjusted p-value < 0.05. Each CpG, which was significant within any of these comparisons, was considered a differentially methylated site (DMS). To validate the numbers of DMS between pairs of castes, we repeated this analysis for 1000 random pairings of 3 samples, sampled without replacement, and recorded the frequency of DMS in each case.

### GO term enrichment of robustly methylated genes

We extracted the unique list of genes which contained CpGs methylated in all 12 samples (Fig. 2A). A GO-term enrichment test was performed on this list of genes with topGO (version 2.34.075) (Alexa et al., 2010), using the classic algorithm. Node size was set to 5, Fisher exact tests were applied, and we only kept GO terms that matched with 2 genes at least and with a and FDR-value < 0.2.

### Alternative splicing

Alternative splicing was estimated for each gene by measuring differential exon expression with the package DEXseq (Li et al., 2015). This pipeline involves first formatting the gff and then extracting exon read counts from sam files. These sam files had been created in a previous study by mapping RNAseq reads to the *M. natalensis* genome (Séité et al., 2022). The DEXseq pipeline was followed at default settings and for each of the four castes compared to the other three castes, we determined genes containing significantly differentially expressed exons (adjusted p-value < 0.05) relative to whole gene expression. These genes were considered putatively alternatively spliced.

Additionally, we assembled a genome-guided transcriptome from RNAseq data (accessions: SAMN17088123-SAMN17088147) (Séité et al., 2022), using the new tuxedo protocol (Pertea et al., 2016). Raw reads were trimmed using Trimmomatic (v0.38) (Bolger et al., 2014) with parameters TRAILING:25 LEADING:25 SLIDINGWINDOW:4:20 AVGQUAL:20 MINLEN:50. Only reads with both pairs after trimming were used for the further analysis. The trimmed RNAseq reads were mapped to the genome with Hisat2 (v2.1.0) (Kim et al., 2019) at default settings for each library. Individual transcriptomes were assembled and merged into one with StringTie (v1.3.4) (Pertea et al., 2016). Numbers of transcripts per annotated gene were then extracted from the resulting gff.

### Differential expression

All data on gene expression levels and caste- and age-biased expression were obtained from Séité et al. (2022).

## Availability of data and material

RRBS sequences have been deposited on NCBI, available under the accession PRJNA742659. Scripts and detailed methods are available on the github repository https://github.com/MCH74/Mna_Methylation.

## Authors’ contributions

M.V.-C. conceived the project and provided biological materials. D.S.D and M.V.-C. collected wild samples. S.G. carried out RRBS services. M.C.H. & E.D. carried out all bioinformatics analyses. M.C.H., S.S. & M.V.-C. interpreted data. M.C.H. wrote the manuscript with contributions from all authors.

## Acknowledgements

We are grateful for valuable guidance and input from Erich Bornberg-Bauer and to Alain Robert for field assistance.

## Supplementary Material

### Supplementary Figures

**Figure S1:**
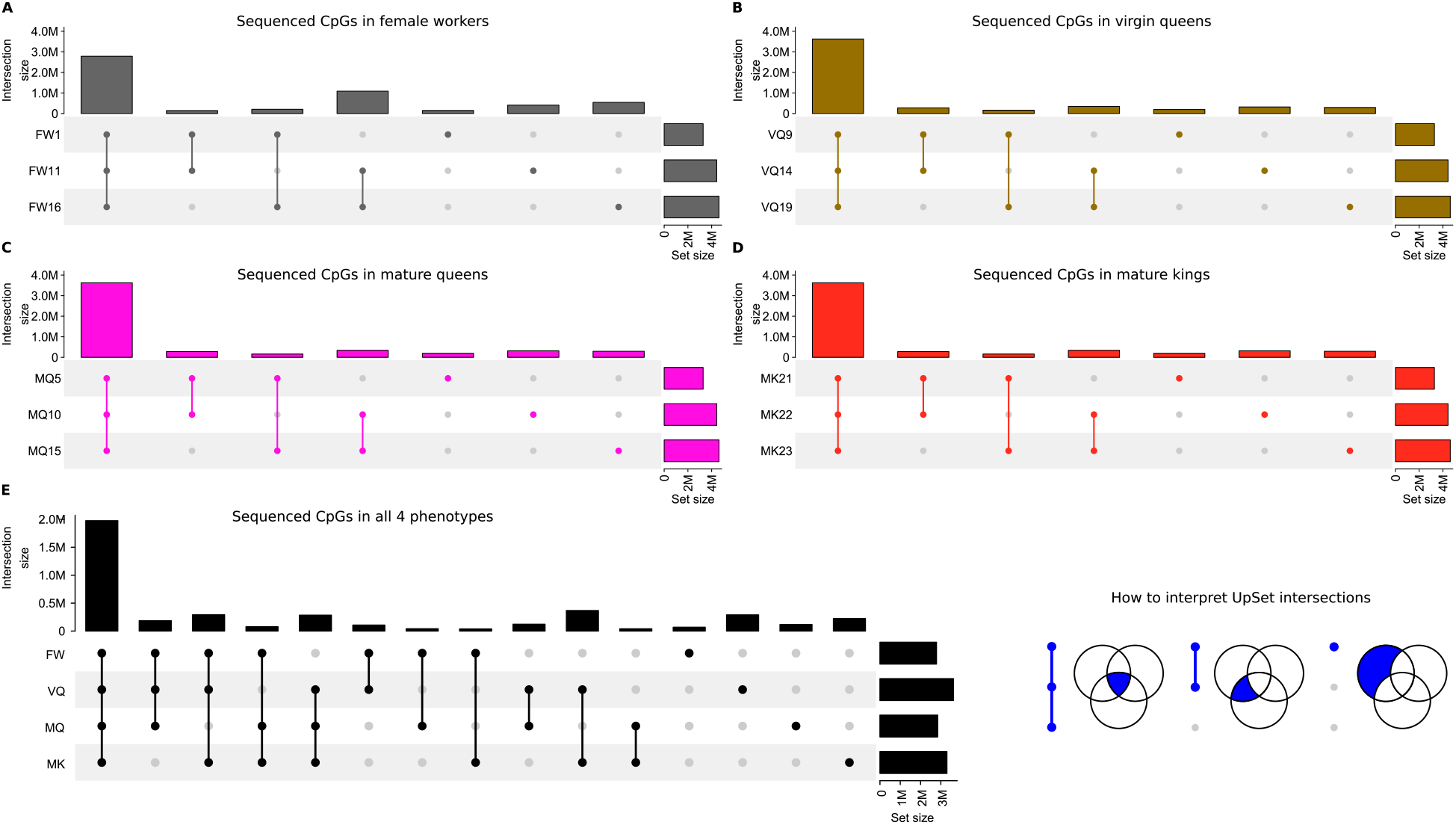
RRBS coverage across 4 phenotypes and 3 replicates. The UpSet plots in visualise sizes of intersections between sets. The central matrix in each shows with joined, coloured dots, which sets are included in the intersections (see inset at bottom right), the vertical columns show the size of these intersections and the horizontal bars show the set sizes. Specifically, **A.-D.** show numbers of sequenced CpGs for each of the three replicates (horizontal bars) and their overlaps between replicates (vertical bars) in female workers (FW), virgin queens (VQ), mature queens (MQ) and mature kings (MK), respectively. **E.** shows total numbers of sequenced CpGs covered by all four phenotypes (FW, VQ, MQ, MK), and how they overlap between phenotypes. In **E.**, sets are comprised of those CpGs which were sequenced in all three replicates represented by left-most vertical bar in plots **A.-D.**.

### Supplementary Tables

**Table S1:** Summary of the sampling design and sequencing results. This table contains the origin of the fat body obtained from the 4 *Macrotermes natalensis* phenotypes (female worker, FW; virgin queens, VQ; mature queens, MQ; mature kings, MK) analysed in this study. These termites were collected from field colonies (colony ID) in 2016 in Southern Africa as described in (Séité et al., 2022). The number of individuals pooled per sample, the bisulfite non-conversion rates, total numbers of sequenced reads, and mapping rates are indicated. The same fat body samples were used to prepare total RNA for transcriptomes (Séité et al., 2022) and for genomic DNA for methylome analyses presented in this manuscript.

| Sample-ID | Phenotype | Colony-ID | Nr. pooled individuals | non-conversion rate | Nr. reads | mapping rate |
| --- | --- | --- | --- | --- | --- | --- |
| 1 | FW | 3 | 85 | 0.9 | 32.1M | 67.3% |
| 11 | FW | 6 | 85 | 0.7 | 47.4M | 68.3% |
| 16 | FW | 7 | 85 | 0.8 | 57.4M | 68.4% |
| 9 | VQ | 5 | 10 | 0.9 | 47.7M | 69.9% |
| 14 | VQ | 6 | 10 | 1.1 | 59.4M | 70.1% |
| 19 | VQ | 7 | 10 | 1.4 | 48.9M | 71.2% |
| 5 | MQ | 3 | 1 | 0.6 | 35.2M | 70.5% |
| 10 | MQ | 5 | 1 | 0.6 | 61.4M | 70.8% |
| 15 | MQ | 6 | 1 | 0.9 | 45.5M | 70.3% |
| 21 | MK | 5 | 1 | 1.1 | 47.3M | 69.8% |
| 22 | MK | 6 | 1 | 0.7 | 51.7M | 70.5% |
| 23 | MK | 7 | 1 | 0.9 | 40.1M | 69.3% |

**Table S2:**
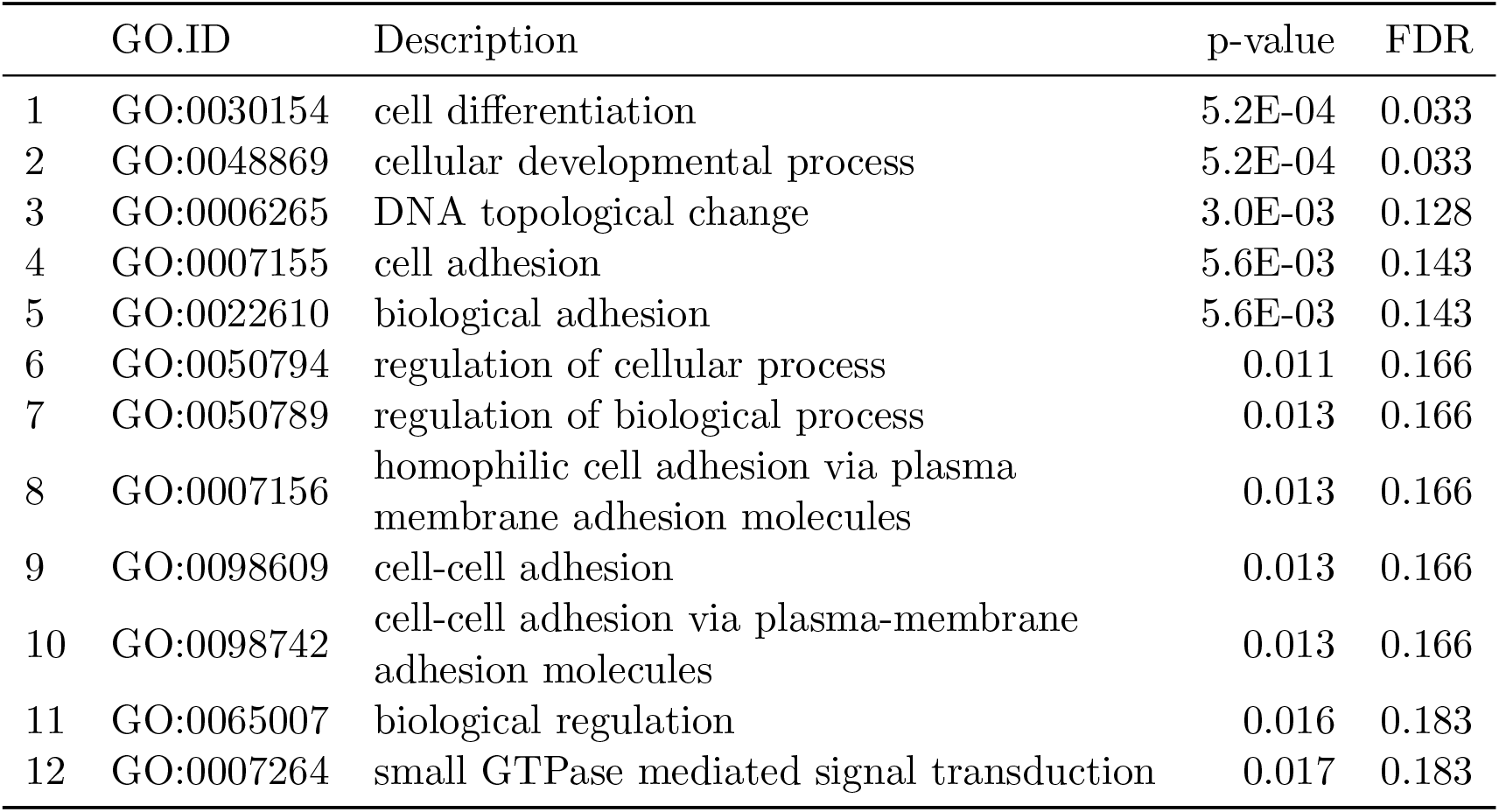
GO-terms significantly enriched among genes containing robustly methylated CpGs, i.e. in all 12 samples. Shown are all terms with an FDR < 0.2.

**Table S3:**
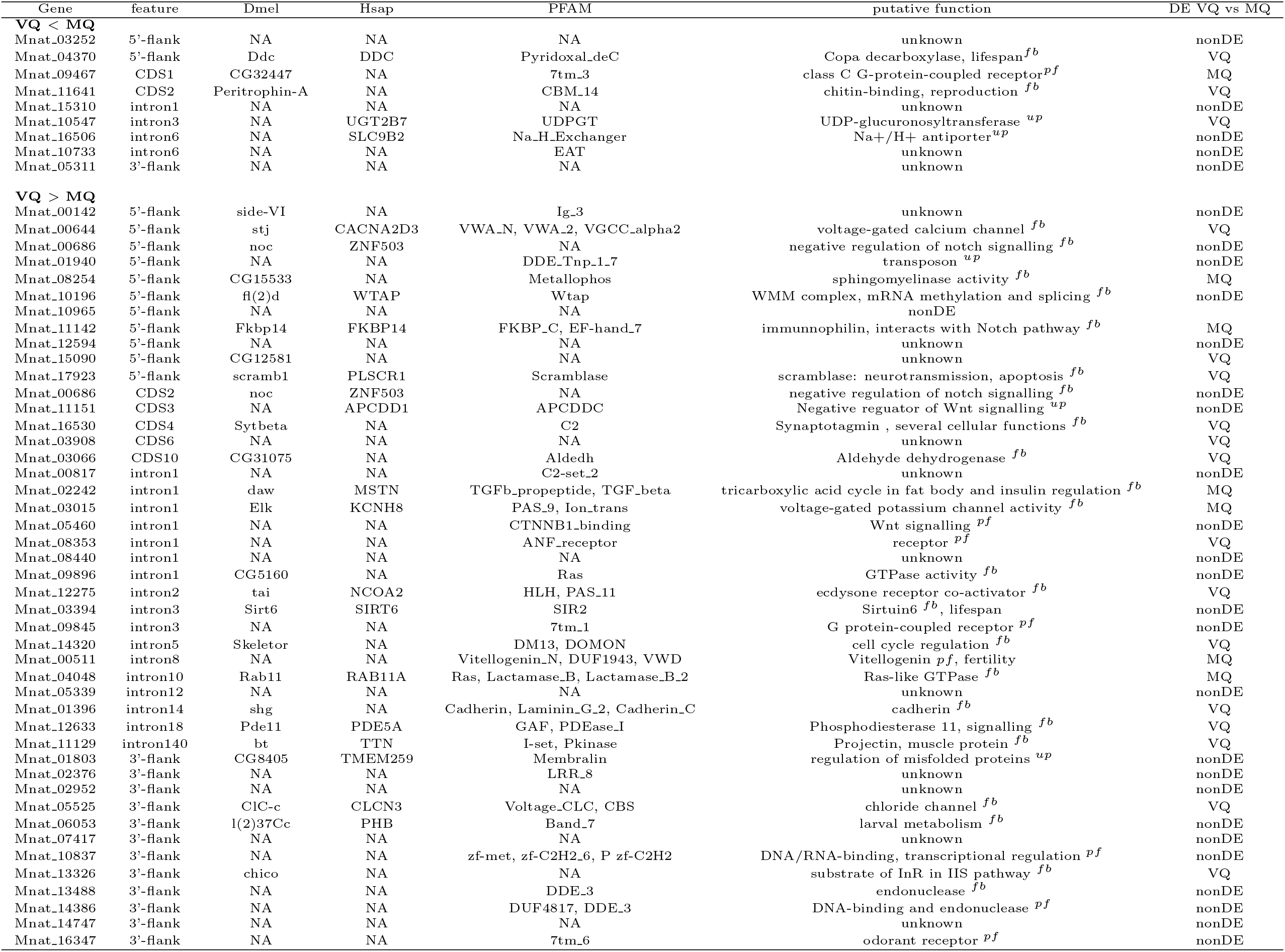
Genes containing significantly differentially methylated sites between young, virgin queens and mature queens. Dmel & Hsap: ortholog in *Drosophila melanogaster* and *Homo sapiens*. DE: differential expression between VQ & MQ (Séité et al., 2022).

